# Dynamic Data Independent Acquisition Mass Spectrometry with Real-Time Retrospective Alignment

**DOI:** 10.1101/2022.11.29.518428

**Authors:** Lilian R Heil, Philip M Remes, Jesse D Canterbury, Ping Yip, William D Barshop, Christine C Wu, Michael J MacCoss

**Author notes:** Corresponding Author Michael J. MacCoss.

## Abstract

We report a data independent acquisition (DIA) strategy that dynamically adjusts the tandem mass spectrometry (MS/MS) windows during the chromatographic separation. The method focuses MS/MS acquisition on the most relevant mass range at each point in time – improving the quantitative sensitivity by increasing the time spent on each DIA window. We demonstrate an improved lower limit of quantification, on average, without sacrificing the number of peptides detected.

## Main

As mass spectrometry (MS) gains popularity for the detection and quantification of proteins, a variety of data acquisition schemes have been developed to accommodate the diverse range of applications. For untargeted experiments, acquisition methods can be characterized as data dependent or data independent. Data independent acquisition (DIA) systematically collects MS/MS spectra given mass-to-charge (*m/z*) range.^3^ While systematic sampling greatly improves reproducibility between runs, there is an inherent trade-off between the total mass range covered and the quality of the results which must be considered when creating new methods. For example, a wider mass range can theoretically sample a greater range of peptides, but comes at a cost to selectivity, sensitivity, and/or quantitative accuracy, depending on the adjustment made (**Figure S1**). An ideal MS method strikes a balance between maintaining sensitivity by increasing the time per scan, selectivity by minimizing isolation width, coverage by maximizing mass range, and speed by minimizing cycle time. Therefore, we aim to have a method that has the sensitivity and specificity of a narrow DIA precursor isolation window while having the peptide coverage of a wide isolation window while maintaining the same cycle time.

In a typical proteomics experiment, peptides are separated prior to analysis by liquid chromatography (LC). Reversed-phase liquid chromatography is the most widely used separation in proteomics and separates peptides by hydrophobicity.^4^ Larger peptides follow a trend of increasing hydrophobicity^5^. Here we capitalize on the relationship between hydrophobicity and m/z by dynamically adjusting the DIA isolation windows during the separation (**Figure 1**). As a proof-of-concept, we chose to use half as many isolation windows acquired for double the time (and resolution) although a number of different parameters could be adjusted, including cutting the cycle time or isolation window width in half. To minimize the effect of shifts in retention time^6^, we align the current acquisition to a run acquired previously so that the optimal DIA windows are adjusted dynamically^7^.

**Figure 1.**
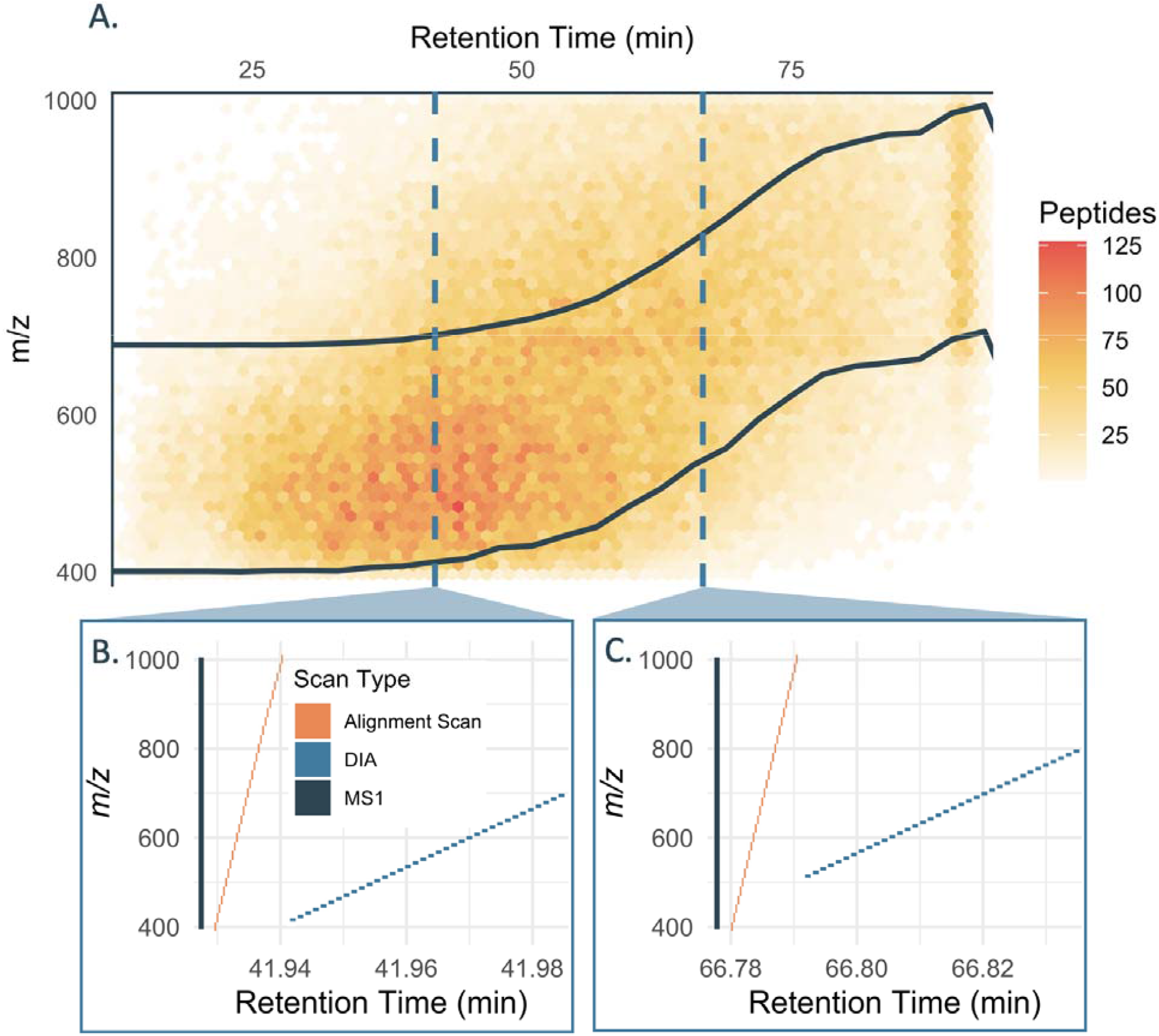
Overview of dynamic DIA method. All features detected in the chromatogram library are binned by retention time and *m/z* with more features corresponding to a lighter color, and the optimal *m/z* boundaries are shown in dark blue (A). The spectra acquired at 41.9 minutes (B) differs from the spectra acquired at 66.8 minutes (C) only in the *m/z* range of precursor windows that were isolated.

To create a dynamic DIA method, we start by creating a chromatogram library using gas phase fractionation DIA (**Methods**).^8^ This chromatogram library serves two purposes: it provides a reduced list of peptides detectable in the quantitative runs^9^ and is used to optimize the isolation window placements for scheduled DIA. The optimal DIA windows at different time points across the gradient are selected to maximize the number of library peptides covered while achieving the desired cycle time (**Figure 1A**). To use dynamic scheduling, we adapt the method described by Remes *et al*. (2020).^7^ Briefly, each instrument cycle consists of an MS1 scan, a set of fast DIA alignment scans covering the entire mass range acquired in the linear ion trap, and a set of high-resolution scheduled DIA spectra (**Figure 1 B&C**). We acquire an alignment run using the scheduling from the original library acquisition. Because this run is acquired soon after the library acquisition, shifts in retention time will be minimal and scheduling is not necessary. The alignment run is preprocessed using Haar compression and subsequent, quantitative runs are acquired using the same sets of DIA alignment scans to determine retention time shifts relative to the alignment run, updating the scheduling accordingly. We have contemplated and tested several improvements to the methodology that don’t change its essence as presented here, for example by including the alignment scans as part of the chromatogram library acquisition to avoid the extra alignment run; or by using “self-alignment” of the DIA scans to eliminate extra alignment acquisitions all-together.

As proof of concept, we applied this dynamic DIA method to a HeLa cell lysate, generating a chromatogram library with 115,917 peptides spanning an *m/z* range of 400-1000. We elected to keep the isolation window width the same as our traditional DIA method, while doubling fill time and resolution. Therefore, we were theoretically able to measure 81,833 of the library peptides (70.6%) (**Figure 1A**). Because we use gas phase fractionation to acquire chromatogram libraries, we expect the quantitative data to contain fewer detectable peptides than the theoretical limit.

We used a matrix-matched calibration curve to compare the quantitative performance of dynamic DIA relative to our gold standard DIA method.^10^ We generated calibration curves and determined the lower limit of quantitation (LoQ) (**Methods)**.^10^ Here, we define peptides as quantifiable if they were able to be assigned a lower limit of quantitation less than the undiluted sample.

With traditional DIA, we were able to detect 34,443 peptides in the matrix-matched calibration curve experiment, of which 23,888 were deemed quantifiable based on their LoQs. With dynamic DIA, we detected 36,355 peptides, and quantified 24,597 (**Figure 2**). Further, the median LoQ is a relative concentration of 20% with traditional DIA compared to 16% for dynamic DIA (**Figure 2B)**. We observe almost a 2 fold improvement in LoQ with dynamic DIA compared to the traditional static method in peptides that can be quantified from both datasets (**Figure 2C**). Because dynamic DIA focuses on a smaller *m/z* range, the instrument spends more time measuring peptides in this range and more ions are measured per spectrum (**Figure S1**).

**Figure 2.**
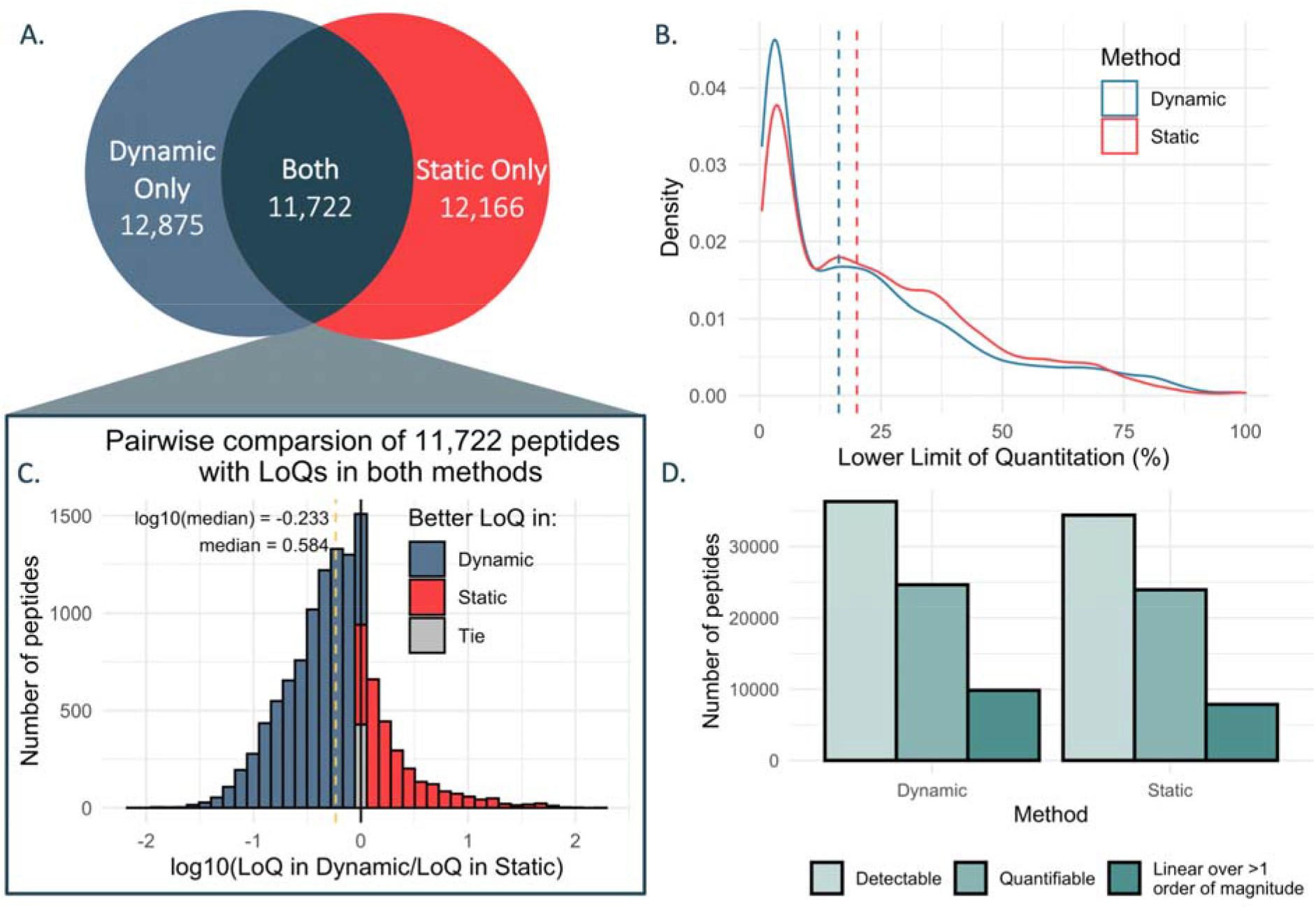
Comparison of quantifiable peptides using dynamic and traditional DIA. The total number of peptides that are deemed quantifiable in each method are shown with overlaps between the methods indicated (A). The relative density of limits of quantitation for all peptides in each method suggests that the dynamic DIA method typically leads to better LOQs (B). In cases where peptides are quantifiable in both methods, there is over a two-fold improvement (reduction) in lower limit of quantitation when using the dynamic method (C). Dynamic DIA detects and quantifies more peptides than traditional DIA (D).

In this work, we have demonstrated the potential of scheduled DIA to improve quantitative sensitivity and precision while maintaining the number of peptide detections. This method is ideally suited to shorter chromatographic gradients with narrow elution peaks, cases where peptides of interest span a large m/z range, and complex mixtures where narrow isolation windows are important for selectivity. Narrow isolation windows could be especially useful when using unit resolution mass analyzers, where interference is a concern. Narrowing acquisition windows is especially important given recent work showing that the unit resolution linear ion trap is more sensitive than Orbitrap analyzers in both targeted and DIA methods.^12,13^ Such sensitivity improvements are critical advancements for advancing any mass spectrometry-based measurements, including the still nascent field of single cell and low input proteomics, an area that has generated widespread interest in recent years.

## Methods

### Cell Culture

HeLa S3 cells were stable isotope labeled with SILAC using the Thermo Scientific SILAC Protein Quantitation Kit with DMEM (Catalog A33972, ThermoFisher Scientific) as recommended by the manufacturer. Briefly, cells were cultured in DMEM/10% FBS and then exchanged into DMEM (for SILAC) containing either ^13^C_6_ ^15^N_2_ L-lysine and ^13^C_6_ ^15^N_4_ L-arginine or unlabeled L-lysine and L-arginine and 10% dialyzed FBS in 80 cm^2^ flasks. Media was also supplemented with Penicillin-Streptomycin and GlutaMAX (Thermo Scientific). Cells were labeled for 8 cell doublings by maintaining log phase growth by exchanging labeled media and splitting as necessary. After washing with PBS, cells from each flask were harvested at ∼85% confluency and pelleted by centrifugation at 1500 x g for 10 min. Cell pellets were stored at -80°C.

### Sample Preparation

The cell lysates were prepared using an automated single pot solid phase sample preparation (SP3) protocol.^14^ Briefly, cell pellets were lysed in 2% SDS with 10 seconds of probe sonication. Total protein concentration was estimated using a Pierce BCA assay (Thermo Fisher Scientific) and 1% SDS solution was added to bring the final concentration to approximately 4 μg/μL. Proteins were reduced in 20 mM dithiothreitol and alkylated in 40 mM iodoacetamide. Following reduction and alkylation, each sample was diluted to 70% acetonitrile and bound to MagResyn Hydroxyl particles (Resyn Biosciences) at a ratio of 100:1 (beads to protein) for a total of 10 minutes.

Subsequent washing and digestion steps were performed on a Kingfisher Flex (Thermo Fisher Scientific). The samples were washed three times in 95% acetonitrile and twice in ethanol. Then, samples were digested with trypsin in 50 mM ammonium bicarbonate at an enzyme to protein ratio of 1:25. The peptide samples were dried down via vacuum centrifugation and resuspended in 0.1% formic acid. A dilution curve was made by diluting the normal digest in the SILAC labeled digest with an equal amount of Pierce Peptide Retention Time Calibration standard (Thermo Fisher Scientific) spiked into each sample. A total of 11 dilution points were made, consisting of the following fractions of normal digest: 0, 0.5, 1, 3, 5, 7, 10, 30, 50, 70, and 100%.

### LC-MS/MS analysis

LC-MS/MS analysis was performed with a Thermo Easy-nanoLC coupled to a Thermo Orbitrap Lumos Tribrid mass spectrometer run in development mode. Peptide separation was performed with an in-house manufactured column consisting of 75 μm inner diameter fused silica capillary (TSP075375, Molex) pulled to an approximately 5 μm tip using a CO2 laser-based micropipette puller (Sutter Instruments). The capillary was packed to a final length of 30 cm with 3 μm ReproSil-Pur C18 beads (Dr. Maisch GmbH) using an in house-built pressure bomb with high pressure helium gas. A trap column was created by packing a 150 μm inner diameter fused silica capillary (TSP150375, Molex) fritted on one end with Kasil with approximately 3 cm of 3 μm ReproSil-Pur C18 beads (Dr. Maisch GmbH). Mobile phase A was 0.1% formic acid in water and mobile phase B was 0.1% formic acid in 80% acetonitrile. Peptide separation was performed via reversed-phase liquid chromatography over a 90 minute gradient from 0 to 40% B followed by a 5 minute ramp to 75% B and a 5 minute hold at 75% B.

Data for the chromatogram library were acquired on the instrument setup prior to quantitative runs. Tandem mass spectra were acquired in 6 gas-phase fractions with 4-*m/z* staggered DIA windows as described in detail by Pino et al. (2020).^8^ After the library was acquired, a retention time alignment run was performed, using the dynamic DIA method described below without any retention time alignment. This run was preprocessed and used to align all future dynamic runs.

For quantitative analysis, two separate DIA methods were used: a traditional static method and a dynamic DIA method. The traditional static DIA method consisted of one MS1 spectrum followed by a set of MS2 spectra acquired at 15k resolving power with 8-*m/z* isolation windows covering a span of 400-1000 *m/z*. The dynamic DIA method began with the same MS1 spectrum covering the full 400-1000 *m/z* range. In addition, it included a set of alignment DIA spectra acquired in the linear ion trap with 20-*m/z* isolation windows. These spectra were used to align the next set of high-resolutionon spectra to a reference run that was acquired immediately after chromatogram library acquisition. This third set of spectra consisted of 8-*m/z* isolation windows at 30k resolving power, covering a variable *m/z* range of approximately 300 *m/z*. Based on the retention time alignment, the exact m/z positioning of these spectra was adjusted over time. The bounds of these spectra were optimized as described below.

The matrix-matched calibration curve was acquired by injecting each dilution point twice, acquiring once with the static DIA method and a second time with the dynamic method. The curve was run from low concentration to high concentration, switching the order of the DIA methods for each point. In total, three replicates of the curve were acquired for both DIA methods.

### Chromatogram library creation

A chromatogram library was created by analyzing the gas phase fractionated DIA data in EncyclopeDIA (v 1.4.10).^9^ Each fraction was searched against a Prosit predicted library^15^ generated from the human Uniprot database. A detailed description of this process is provided by Pino et al. (2020).^8^ The resulting chromatogram library was exported for downstream analyses.

### Selection of dynamic DIA windows

To optimize the window boundaries for the dynamic DIA window, a maximization method was used. The code to do so is available on GitHub, and the process is described conceptually here. First, the window span was determined based on the instrument acquisition rate, desired isolation width, and desired points across the peak. At 30k resolving power, the instrument acquisition rate is 14.6 Hz. To acquire 10 points across LC peaks that are on average 25 seconds wide at the base requires a cycle time of 2500 msec; therefore, there is time for at most 2.5 s * 14.6 Hz = 36 acquisitions per cycle. Using 8-*m/z* isolation windows means that a 300-*m/z* span can be measured with these 36 acquisitions. All of the features in the chromatogram library were binned by retention time and *m/z*. For each time bin, the *m/z* range that covers the most detectable features from the chromatogram library was selected. These window boundaries with scheduling were uploaded directly into the instrument method for the dynamic DIA setting.

### Real time alignment

Real time retrospective retention time alignment was performed by the instrument as described by Remes et al. (2020).^7^ Briefly, the retention time shift relative to a reference run is determined based on the average cross correlation of a set of high-speed DIA spectra acquired in the linear ion trap in a reference run relative to the current run. The retention time shift is used to determine the active targets and acquire data for these targets. In this case, rather than specific scheduled peptide targets, scheduled DIA windows were used. The DIA spectra were listed as targeted MS2 spectra with specific retention time windows, thus the execution of this method is the same as described in the original publication.

### Data processing

Data were searched with EncyclopeDIA (v 1.2.2)^9^ using the protocol described by Pino et al. (2020).^8^ All quantitative files were searched against the sample-specific chromatogram library and filtered for peptides at 1% FDR with at least three quantitative transitions to generate a new quantitative chromatogram library. Using the quantitative library for each method respectively, quantitation was performed in Skyline^11^, and the areas under the curve for each peptide were used for quantitation. Lower limits of detection and quantitation were determined using a bootstrapping method.^10^ Further analysis of the results was performed in R.

## Data Availability

Raw data files, EncyclopeDIA search results, and all Skyline documents have been deposited in the ProteomeXChange Consortium via Panorama Public (https://panoramaweb.org/dynamicDIA.url).

## Code Availability

Analysis code is available on GitHub,https://github.com/uw-maccosslab/dynamic-DIA.

## Acknowledgements

This work was supported in part by National Institutes of Health grants U19 AG065156 and R24 GM141156.

## Notes

The authors declare the following competing financial interest(s): The MacCoss Lab at the University of Washington has a sponsored research agreement with Thermo Fisher Scientific, the manufacturer of the instrumentation used in this research. M.J.M. is a paid consultant for Thermo Fisher Scientific. P.M.R., J.D.C., P.Y., and W.D.B., are employees of Thermo Fisher Scientific, the manufacturer of the instrumentation used in this research.

## Supplementary information

Supplemental File S1: Supplemental figures referenced in text

