## Supplemental File S1 for "Dynamic Data Independent Acquisition Mass Spectrometry with Real-Time Retrospective Alignment"

### Supporting information for: Dynamic Data Independent Acquisition Mass Spectrometry with Real-Time Retrospective Alignment

Lilian R Heil[1], Philip M Remes[2], Jesse D Canterbury[2], Ping Yip[2], William D Barshop[2], Christine C Wu[1], Michael J MacCoss[1*]

[1] Department of Genome Sciences, University of Washington, 3720 15th Street NE, Seattle, Washington 98195, United States

[2] Thermo Fisher Scientific, 355 River Oaks Parkway, San Jose, California 95134, United States

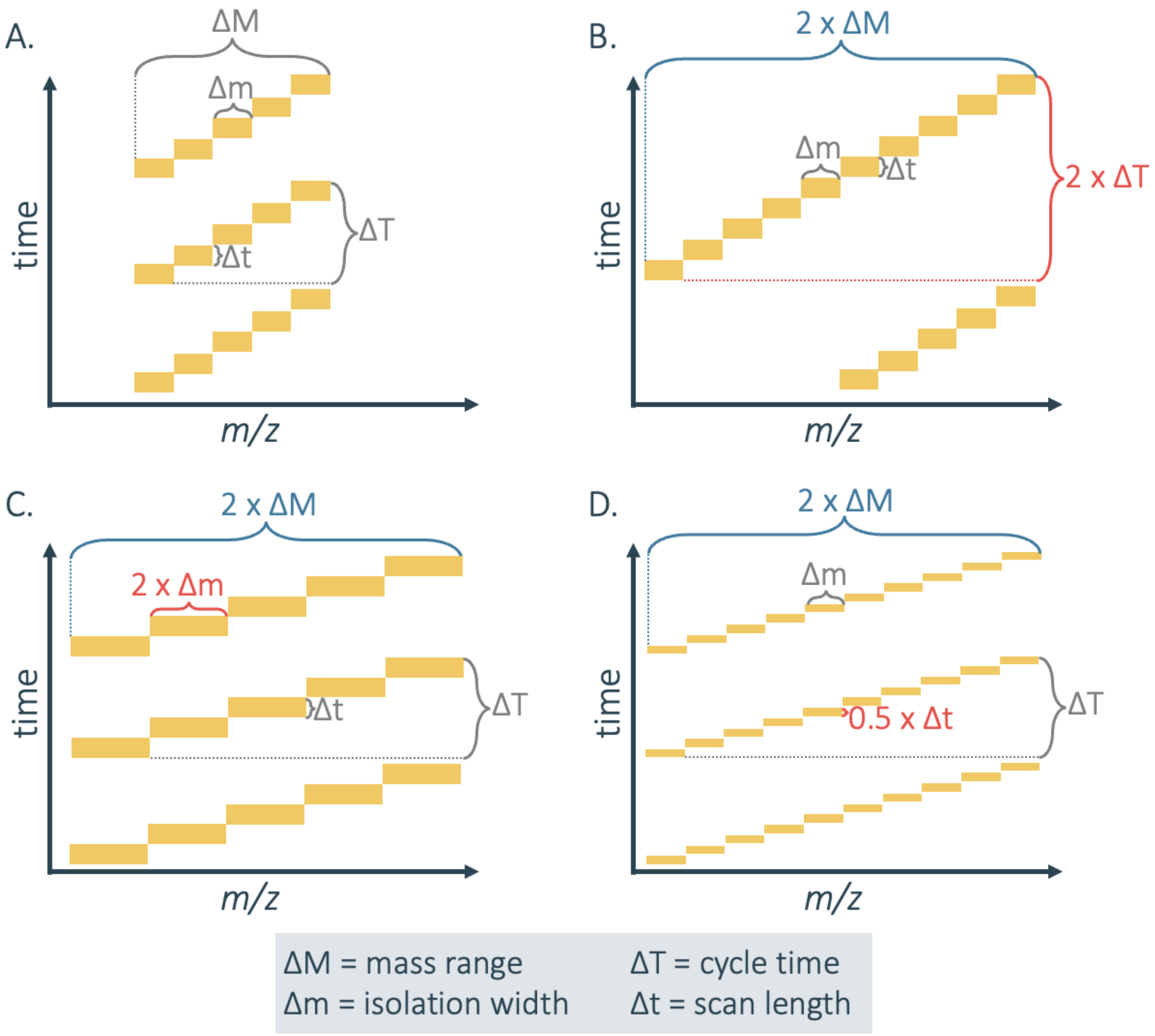


**Figure S1.** *Tradeoffs associated with DIA methods*. A wider mass range can theoretically sample a greater range of peptides, but comes at a cost to selectivity, sensitivity, and/or quantitative accuracy, the exact nature of which depends on the isolation width selected. A generic DIA method is shown in (A), with (B-D) illustrating possible alterations to compensate for doubling the mass range. The parameter negatively impacted by doubling the mass range is shown in red. If the isolation width is kept narrow, the amount of time that can be spent acquiring each spectrum and the number of data points acquired per LC peak is limited, decreasing sensitivity, accuracy, and reproducibility (B). If the isolation width is increased, selectivity, dynamic range, and sensitivity may suffer (C). On the other hand, a narrower mass range improves data quality by allowing a combination of either improved selectivity or sensitivity, at the cost of reduced peptide coverage (D).

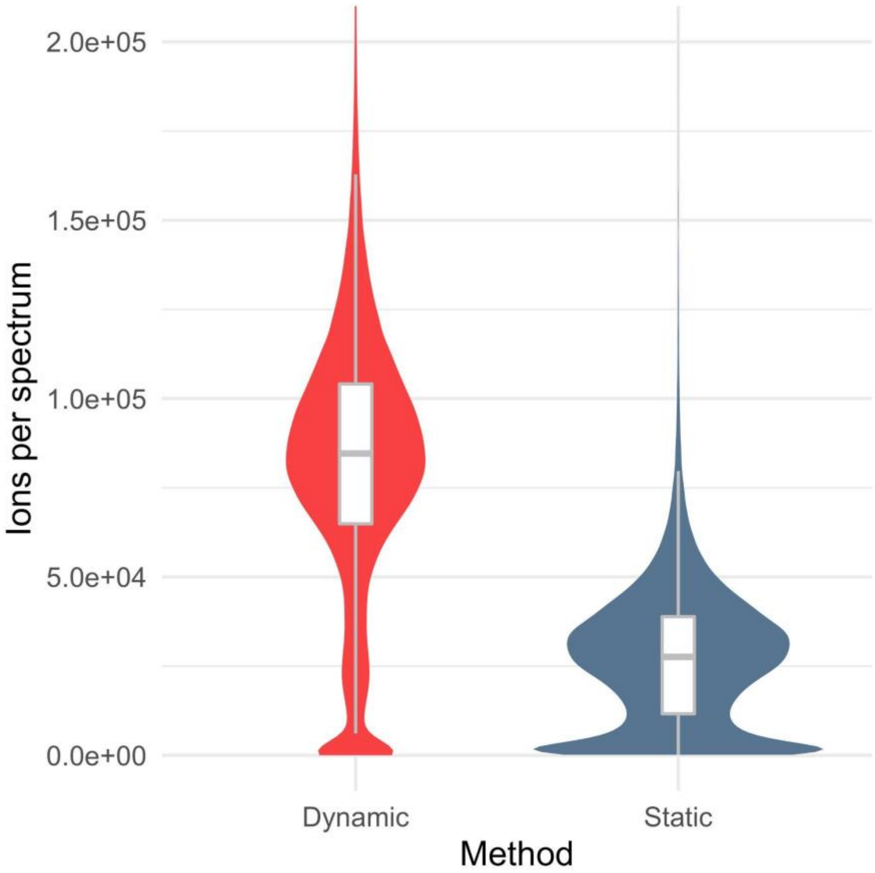


**Figure S2.** *Number of ions per spectrum*. By leveraging the dynamic DIA method to increase maximum injection times allowed by automatic gain control, each spectrum contains more ions which accounts for the increased sensitivity.
